# Pathfinder: open source software for analyzing spatial navigation search strategies

**DOI:** 10.1101/715961

**Authors:** Matthew B. Cooke, Timothy P. O’Leary, Phelan Harris, Richard E. Brown, Jason S. Snyder

**Affiliations:** Department of Psychology, Djavad Mowafaghian Centre for Brain Health, University of British Columbia, 2211 Wesbrook Mall, Vancouver, British Columbia, Canada V6T 2B5; Psychology and Neuroscience Department, Dalhousie University, Halifax, Nova Scotia, Canada B3H 4R2

**Author notes:** Author Contributions: Developed software (MBC, PH), performed experiments (TPO), analyzed data (MBC, TPO, JSS), wrote manuscript (JSS), edited manuscript (MBC, TPO, REB), provided funding (JSS, REB).

**Keywords:** search strategy, water maze, learning, memory, rodent, reversal, goal

## Abstract

Spatial navigation is a universal behavior that varies depending on goals, experience and available sensory stimuli. Spatial navigational tasks are routinely used to study learning, memory and goal-directed behavior, in both animals and humans. One popular paradigm for testing spatial memory is the Morris water maze, where subjects learn the location of a hidden platform that offers escape from a pool of water. Researchers typically express learning as a function of the latency to escape, though this reveals little about the underlying navigational strategies. Recently, a number of studies have begun to classify water maze search strategies in order to clarify the precise spatial and mnemonic functions of different brain regions, and to identify which aspects of spatial memory are disrupted in disease models. However, despite their usefulness, strategy analyses have not been widely adopted due to the lack of software to automate analyses. To address this need we developed Pathfinder, an open source application for analyzing spatial navigation behaviors. In a representative dataset, we show that Pathfinder effectively characterizes the development of highly-specific spatial search strategies as male and female mice learn a standard spatial water maze. Pathfinder can read data files from commercially- and freely-available software packages, is optimized for classifying search strategies in water maze paradigms, but can also be used to analyze 2D navigation by other species, and in other tasks, as long as timestamped xy coordinates are available. Pathfinder is simple to use, can automatically determine pool and platform geometry, generates heat maps, analyzes navigation with respect to multiple goal locations, and can be updated to accommodate future developments in spatial behavioral analyses. Given these features, Pathfinder may be a useful tool for studying how navigational strategies are regulated by the environment, depend on specific neural circuits, and are altered by pathology.

## INTRODUCTION

All living organisms move throughout space to survive. Amongst mammals, there is a diversity of spatial behaviors that depend on numerous factors such as anxiety^1,2^, learning^3^, and the nature and pattern of stimuli that predict goals^4–6^. Given rodents’ natural propensity to explore stimuli and environments, an array of rodent navigational tasks have been developed to investigate how various brain regions interact to control goal-direct behavior^7^. This has routinely been conducted using fixed-trajectory mazes such as the T-maze or radial maze. While these dry maze paradigms offer the convenience of fixed choice points that reduce ambiguity associated with classifying decisions and navigational responses, they cannot be used to study patterns of exploration in open environments.

A popular approach for studying free navigation in animals has been the water maze, where rodents learn the location of a hidden escape platform in a pool of water based on distal and/or local cue configurations^3^1. Early studies validated the usefulness of the water maze for studying spatial processing and described progressive stages of learning where a rodent searches for the platform with increasing spatial specificity^8,9^. The vast majority of studies have since used escape latency or path length as primary measures of spatial learning. However, water maze navigation is unconstrained and animals can solve the task using different strategies that may not always differ in terms of the time it takes to reach the platform^8,9^. Thus, while latency and path length measures are convenient, they discard a rich amount of behavioral data.

Over the years, a number of groups have described manual and automated methods for classifying search strategies used by animals and humans in water maze experiments^8,10–19^. By mathematically relating the swim path to features of the maze environment one can identify and quantify the types of search strategies employed. Search strategy analyses have revealed that the ventral hippocampus is involved in coarse spatial goal-directed search^16^, that adult neurogenesis promotes spatially precise search^20^, and that spatially accurate search is reduced in humans with, and/or animal models of, Alzheimer’s disease^21,22^, autism^23^, traumatic brain injury^22,24^ and aging^14,25^. Despite the utility of these analyses they have been relatively uncommon to date, likely because commercially-available software packages often do not perform these analyses and the analytic methods used in previous work are not typically available in the form of an easy-to-use software package.

To facilitate the study of navigational search strategies, whether in the water maze or other 2-dimensional navigational paradigms, we created a new software application called Pathfinder. Pathfinder is a Python-based, open source tool with an intuitive graphical user interface and adjustable parameters for conducting detailed analyses of spatial search patterns. We validate Pathfinder with a mouse water maze dataset, where we find that male and female mice develop increasingly specific and direct spatial search strategies with additional days of training.

## METHODS

### Installation and Dependencies

Pathfinder is freely available under the GNU General Public License version 3.0. Detailed instructions on use and installation of the program can be found on Github at github.com/MatthewBCooke/Pathfinder. We recommend installing Anaconda for Python 3, as it includes all of the following packages that are needed to run Pathfinder: PIL (https://pillow.readthedocs.io/en/latest/), xlrd (https://xlrd.readthedocs.io/en/latest/), numpy (https://www.numpy.org), pickle (https://docs.python.org/3/library/pickle.html), scipy (https://www.scipy.org), matplotlib (https://matplotlib.org), tkinter (https://wiki.python.org/moin/TkInter). The MATLAB engine is optional and needs to be installed separately for entropy calculations (MATLAB and Statistics Toolbox Release 2018b, The MathWorks, Inc., Natick, Massachusetts, United States). Once Anaconda is installed, Pathfinder can be downloaded via Github or by typing “pip install pathfinder” in a shell window (i.e. Mac terminal or Windows command line). Pathfinder is then opened by typing “pathfinder” into the shell window and pressing return.

### General Usage

Pathfinder has a simple, user-friendly interface for extracting information from spatial navigation tracking files that contain xy coordinates over time (Fig. 1). While it can be used to analyze multiple types of 2D navigational data, it is optimized for rodent spatial water maze experiments and accepts inputs from commonly-used commercial tracking software, including Ethovison (Noldus), Anymaze (Stoelting) and WaterMaze (Actimetrics). Pathfinder can also open files exported from the open source tracking software, ezTrack^26^, enabling a cost-effective and fully open source workflow for detailed water maze behavioral analyses. Trial information from these programs are outputted in CSV or Excel format, which can then be inputted into Pathfinder through the File menu. The experimental setup is specified in the main window (Fig. 1a). Pathfinder can automatically calculate the position and size of the maze and the goal location (provided they are constant across trials), or these parameters can be entered manually.

**Figure 1:**
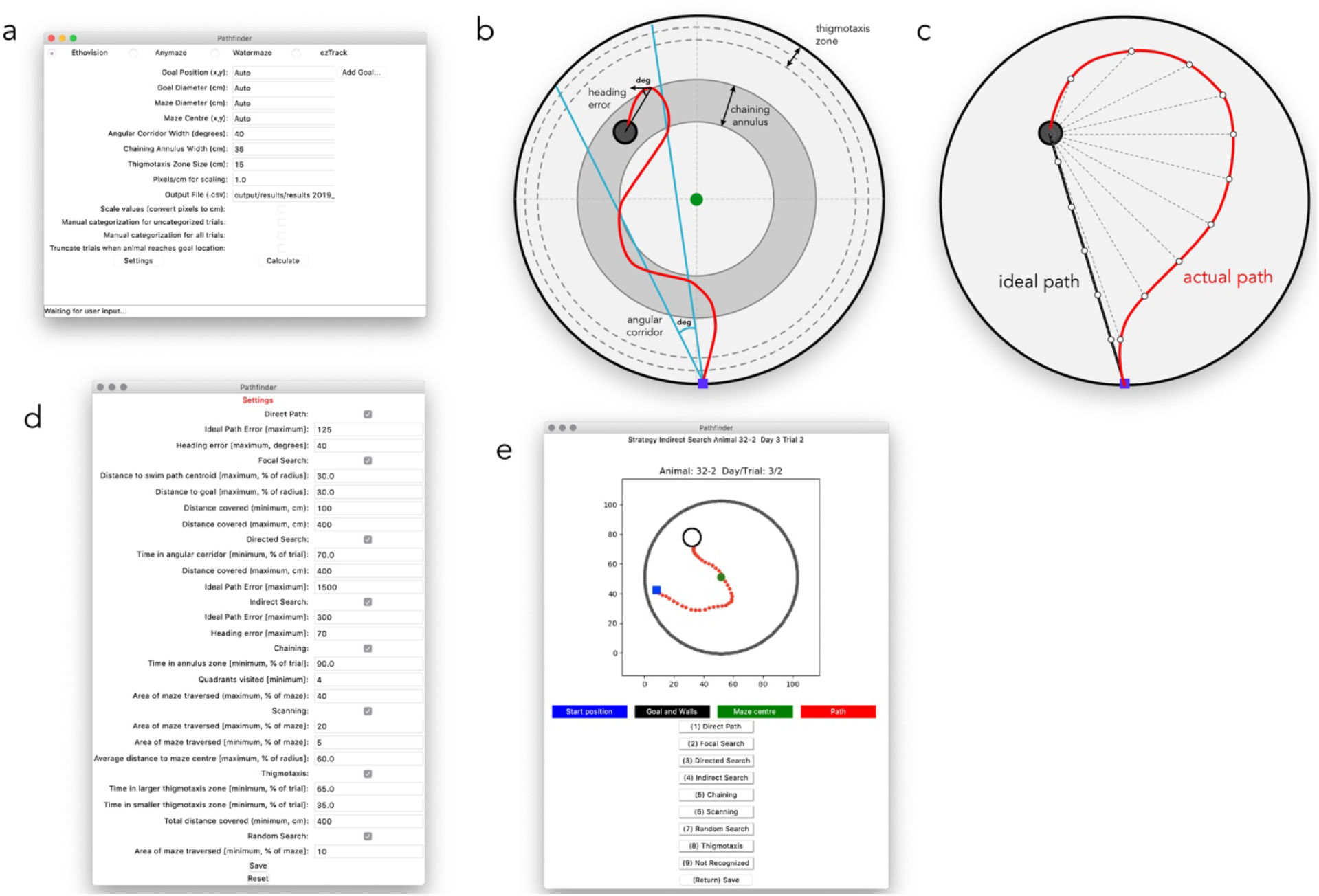
Graphical user interface and setting parameters. a) Screenshot of the main application window, were maze geometry is defined and input and output settings are established. b) Maze schematic and geometry for defining variables. The chaining corridor is centered on the goal platform and extends throughout all 4 quadrants; its width is specified in the main window. The larger thigmotaxis zone is specified in the main window; Pathfinder calculates the smaller thigmotaxis zone as half the width. Heading error is the angular distance between the actual path direction and a straight line to the goal (Pathfinder calculates average heading error at all points; only a single example shown). The angular corridor is used to define the directed search strategy, which depends on the accuracy of the animal’s trajectory as it approaches the platform. The width of the corridor (in degrees) is specified in the main window, and is centered on the goal. c) Schematic of the Ideal Path Error (IPE) metric. The distance from the platform is measured at each timepoint provided by the tracking software (actual path; only a fraction of distances shown for clarity) to provide a cumulative distance measure. Assuming the same swim speed as the actual path, distances are similarly summed from the ideal path, to provide a cumulative ideal path measure. The ideal cumulative distance is subtracted from the actual cumulative distance to generate the IPE. d) Parameter bounds are entered in the settings window. e) The manual categorization window, for viewing trial paths and manually categorizing strategies.

Pathfinder relies on several variables that describe navigation relative to the pool and platform geometry: 1) Ideal Path Error (IPE) is the summed error of the search path (Fig. 1c). It is conceptually similar to the Cumulative Search Error (CSE) since it also measures proximity to the goal throughout the trial^9,27^. An advantage of proximity measures is that they can distinguish 2 trials that have equivalent latencies/path lengths but differ in average distance to the platform. When calculating the IPE, the distance from the goal is measured at each time point in the trial and summed to generate a cumulative distance measure of the actual path (similar to CSE). In contrast to the CSE, the IPE is calculated by subtracting the cumulative ideal path distance from the cumulative actual path distance. The cumulative ideal path is simply the sum of all of the distances between the goal and the position of the animal if it swam along a straight line to escape, using the average velocity from the trial. 2) Heading error is the angular distance between the current path and a straight line to the goal location. The current path direction is defined by a line connecting 2 temporally-adjacent xy coordinates. The average heading error is an average of all of the heading error values for the trial and the initial heading error is the average of the heading error values for the first second of the trial. Additional variables are user-defined on the main window: 3) Angular Corridor Width: the size of the angular navigational corridor (in degrees) that extends from the start location and widens towards the goal, centered on the goal location, 4) Chaining Annulus Width: the width of the chaining annulus, a donut-shaped zone that is centered on the goal and spans all areas of the maze at a fixed distance from the maze wall. 5) Thigmotaxis zone size: the width of a zone that spans the perimeter of the maze and extends inward from the maze wall. Pathfinder also defines a “small” thigmotaxic zone that is half the width of this value. 6) Add goal: Pathfinder will perform all calculations and strategy analyses with respect to an unlimited number of goal locations. This can be used to measure performance and characterize strategies with respect to multiple goal locations (e.g. during spatial reversal, spatial choice). Selecting “truncate trials” will artificially end the trials if/when the subject reaches the additional goal locations. This is necessary, for example, to measure direct trajectories to a former goal location in a reversal paradigm (since the strategy will no longer meet direct path criteria if the former location in contacted and search continues elsewhere in the maze).

Once the variables are defined, boundaries must be set to establish the criteria for strategy categorization. Clicking “settings” will open up an additional window where strategy options can be selected and parameter bounds can be set (Fig. 1d). Upon clicking “calculate”, Pathfinder categorizes trials into one of eight search strategies that are ordered according to the degree of spatial specificity (high to low): 1) direct path, 2) focal search, 3) directed search, 4) indirect search, 5) chaining, 6) scanning, 7) random search, and 8) thigmotaxis. These categories are mutually exclusive and follow a defined order (1 to 8), but the user can opt to exclude strategies from the analysis. Thus, Pathfinder determines, in a stepwise fashion, whether a given trial fulfills the criteria for direct swim. If so, it moves on to categorize the next trial. If not, it determines whether the trial fits the subsequent strategy, and so on. The strategies and their parameters are shown in Fig. 2. In the output file (.csv), each trial is categorized and the following additional metrics are provided: latency and distance travelled to reach the goal, average distance from the goal, percent of maze traversed, velocity, initial and average heading error and IPE. Pathfinder also has the ability to calculate the entropy for each trial, a measure of disorder in the path, relative to the goal location. The entropy calculation calls the MATLAB engine and requires a MATLAB license. Entropy measures the performance by looking at a shift from more disordered swimming (high entropy) to more spatially strategic paths (low entropy), and has been previously found to be highly sensitive to water maze search performance^28^. Due to the manipulation of large matrices, calculating the entropy of trials is very slow.

**Figure 2:**
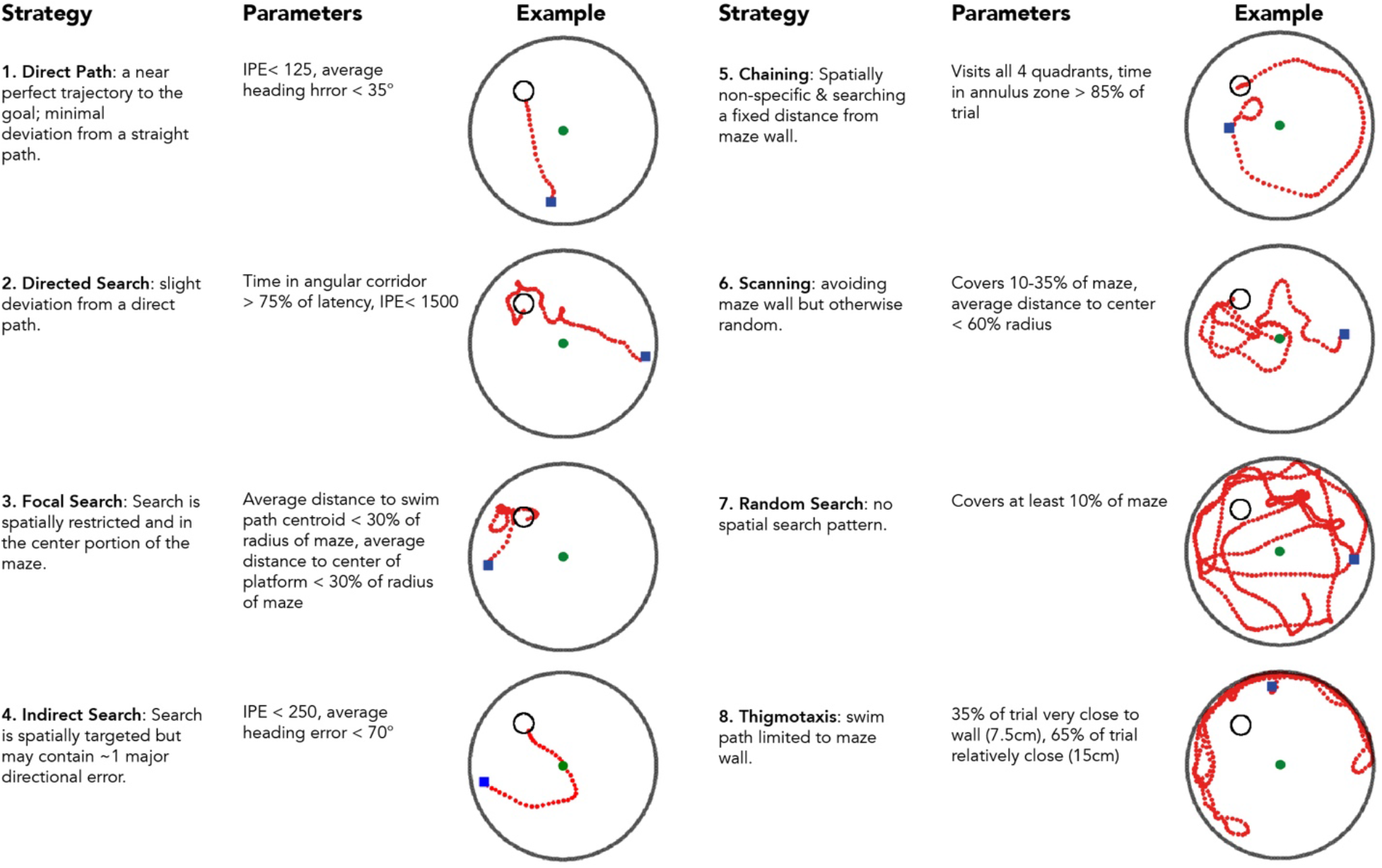
Search strategies and associated parameters. Pathfinder categorizes each trial according to 1 of 8 possible strategies. Categorization proceeds sequentially in the order shown (unless some strategies are excluded from the analysis). For example, for a trial to be classified as Random Search, the path must cover a minimum proportion of the maze *and* not fit any of the criteria for strategies 1-6. In the examples shown, the blue square indicates the start point and the green circle indicates the middle of the pool. Parameter settings are those used in the present study, and should be adjusted depending on changes to testing procedures and maze geometry.

Occasionally, some trials cannot be categorized. The user therefore has the option to manually categorize uncategorized trials, by selecting this option on the main window. Additionally, there is an option to manually categorize all trials. Here, Pathfinder provides an image of the trial as well as shortcut keys to select the appropriate strategy. The software will also display the strategy it had automatically categorized for the displayed trial. Manual categorization will not overwrite the automatic categorization but will be displayed separately in the output file. This allows for comparison between the automatically calculated and user-selected strategy.

In addition to strategy categorization, Pathfinder will also create heatmaps as a useful visual representation of groups of trials. This is accomplished by counting the number of times animal(s) visit each bin in a hexagonal array that is overlaid on the maze (bin size is user-defined). The range of colors (cool to warm) can be automatically set to occupy the full scale. Alternatively, the user can manually set the maximum, above which all bins will read the hottest.

### Animals

A group of 35 C57BL/6J mice (18 male, 17 female) were used in this experiment. Mice were housed in same-sex groups (2-4/cage) in polyethylene cages (30cm x 19 cm x 13cm) with pine chip bedding and a small tube and food and water available *ad-libitum.* Mice were housed under a reversed light-dark cycle (lights off 8:00am-8:00pm), and completed water-maze testing in the dark phase. Mice were first tested on the Barnes maze^29^ and were 18 weeks old when tested on the water maze for the current experiment. All procedures adhered to guidelines from the Canadian Council on Animal Care and were approved by the Dalhousie University Committee on Laboratory Animals.

### Spatial water maze training

The water maze consisted of a plastic circular pool (110 cm diameter) painted black. The pool was filled with water (21-23°C), which was made opaque with the addition of non-toxic white tempera paint (Schola). A circular escape platform (14 cm height, 9 cm diameter) was positioned 1 cm below the water. The water maze was placed in a diffusely lit room with many extra-maze visual cues (posters on walls, a desk, the experimenter, geometric layout of testing room etc.).

Animals were tested over a total of 15 days. They first completed 8 days of acquisition training (A1-A8) with a hidden escape platform (4 trials/day). Across trials, mice were released into the pool from four different locations, with the order differing across mice. They were given a maximum of 60 sec to locate the escape platform, after which they were guided to the platform by the experimenter. Mice remained on the platform for 15-20 seconds before being removed from the pool. During daily test sessions, mice were tested in squads of 4 and each mouse was held in separate cages filled with a bedding of paper towel. The inter-trial interval ranged from 2-8 minutes. The day following acquisition training, memory was assessed with a single 60-sec probe trial with no escape platform present. Mice then completed a single day of re-training (Retrain) to reduce extinction that may occur during the probe trial. During re-training the escape platform is returned to the same location used in acquisition training.

After acquisition re-training, reversal learning was assessed over 3 days (R1-R3) with the escape platform moved to the opposite side of the maze. A reversal probe trial (R probe) was then completed to assess memory for the location of the new escape platform location. Finally, a single day of visible platform training (Visible platform; 4 trials) was completed, where the escape platform was moved to a new location, and made visible with the addition of a striped flag. Behaviour was recorded with the WaterMaze (Actimetrics) video tracking system (5 samples per second), via a camera placed directly above the pool.

## RESULTS

To validate Pathfinder, we trained mice for 8 days on a spatial water maze such that they achieved asymptotic performance according to standard metrics, and should therefore have adopted distinct navigational strategies as they learned the procedural and spatial task demands. Following acquisition, mice received an unreinforced probe trial, 1 day of retraining, 3 days of reversal training (platform in opposite side of pool), another probe trial, and one day of visible platform training (outlined in Fig. 3a).

To confirm that mice learned the task, we first analyzed performance using several metrics that indicate learning but do not reveal details about navigational strategies (Fig. 3). We focussed on acquisition and reversal phases since they are the main focus of our subsequent strategy analyses. Over the 8 days of acquisition, mice reached the platform faster, increasingly swam in the direction of the platform as measured by heading angle error, and had lower IPE and entropy scores. The greatest performance improvements occurred during the first 4 days and, while all measures revealed improvements beyond day 4, only average heading error and entropy analyses revealed improvements beyond day 5. There were no sex differences in acquisition performance.

**Figure 3:**
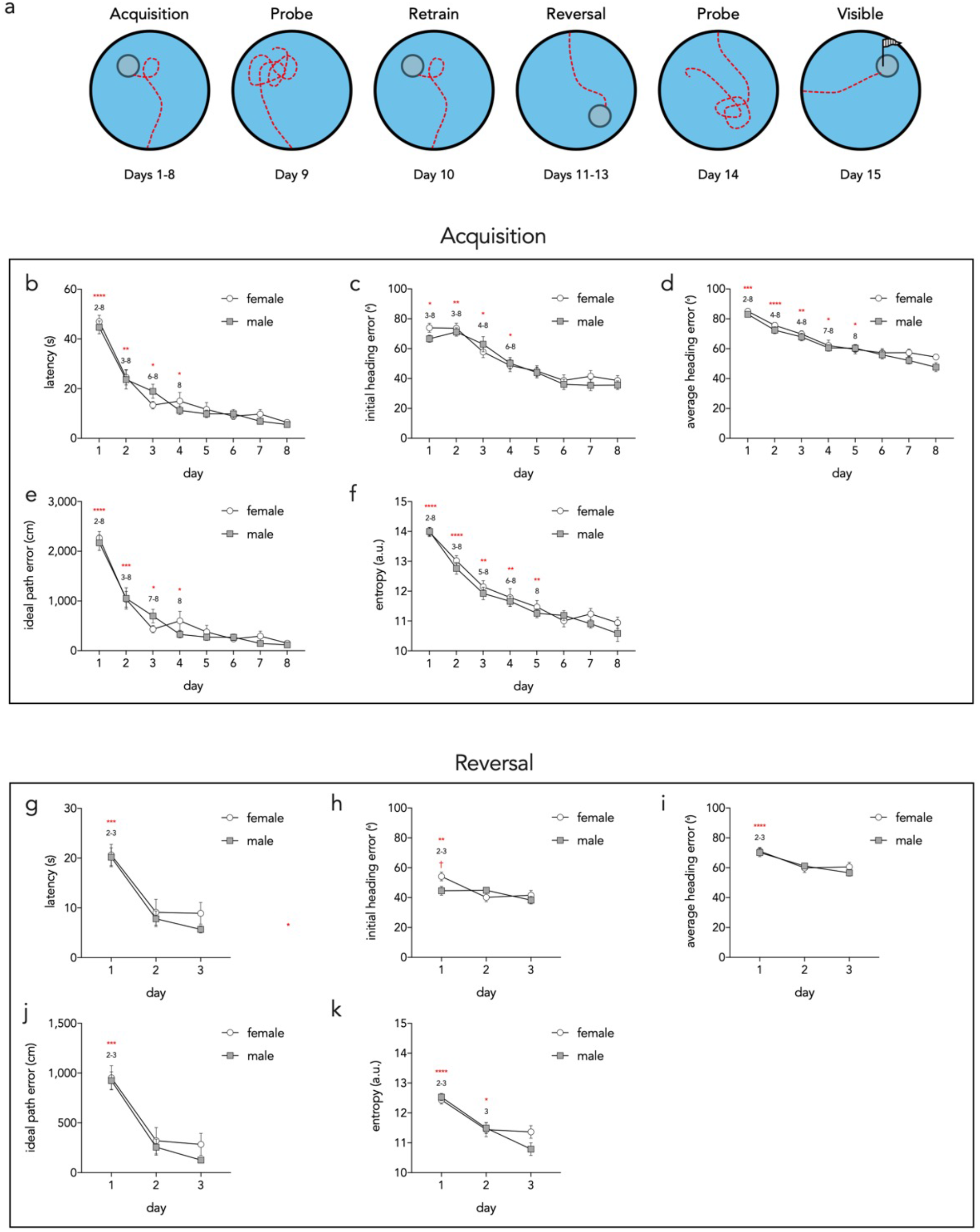
Acquisition and reversal performance as assessed by individual parameters. a) Schematic outline of full behavioral paradigm. Individual performance metrics were analyzed for acquisition (b-f) and reversal (g-k) stages of testing. b) Latency to reach the platform decreased across days (day effect F_7,231_=75, P<0.0001; sex effect F_1,33_=0.3, P=0.6; interaction F_7,231_=0.9, P=0.5). Asterisks denote statistically significant differences from the subsequent days that are indicated by the numbers. c) Initial heading error decreased over days (day effect F_7,231_=39, P<0.0001; sex effect F_1,33_=0.4, P=0.6; interaction F_7,231_=0.7, P=0.6). d) Average heading error decreased over days (day effect F_7,231_=48, P<0.0001; sex effect F_1,33_=2.5, P=0.12; interaction F_7,231_=0.6, P=0.8). e) Idea path error decreased over days (day effect F_7,231_=79, P<0.0001; sex effect F_1,33_=0.3, P=0.6; interaction F_7,231_=1.0, P=0.4). f) Entropy decreased over days (day effect F_7,231_=75, P<0.0001; sex effect F_1,33_=1.3, P=0.3; interaction F_7,231_=0.5, P=0.8). g) Latency decreased over days (day effect F_2,66_=69, P<0.0001; sex effect F_1,33_=0.5, P=0.5; interaction F_2,66_=0.7, P=0.5). h) Initial heading error decreased over days and was greater in females on day 1 (day effect F_2,66_=9, P<0.001; sex effect F_1,33_=1.0, P=0.3; interaction F_2,66_=4.7, P=0.01). i) Average heading error decreased over days (day effect F_2,66_=21, P<0.0001; sex effect F_1,33_=0.2, P=0.7; interaction F_2,66_=0.9, P=0.4). j) Ideal path error decreased over days (day effect F_2,66_=98, P<0.0001; sex effect F_1,33_=0.5, P=0.5; interaction F_2,66_=0.7, P=0.5). k) Entropy decreased over days (day effect F_2,66_=39, P<0.0001; sex effect F_1,33_=0.6, P=0.4; interaction F_2,66_=2.6, P=0.08). *P<0.05, **P<0.01, ***P<0.001, ****P<0.0001, ^†^P<0.05 within day, male vs female comparison. Symbols = mean ± standard error.

Reversal learning performance improvements were mostly apparent after the first day of training, likely because mice had learned the procedural aspects of the task and the spatial environment, and only had to learn a new platform location (Fig. 3g-k)^30^. Path entropy decreased from days 2-3, indicating continued learning. Females and males were equivalent in all performance measures except males had a lower initial heading error on day 1 of reversal training (Fig. 3h).

Pathfinder revealed clear differences in search strategies over days of training (Fig. 4). Over the first 2 days of acquisition, mice were initially thigmotaxic. After learning that the pool wall did not afford escape, they then transitioned to chaining, random and scanning search patterns, all of which indicate spatially non-specific search away from the pool wall. Over days 2-3 mice transitioned to spatially-specific forms of search, with ~30% performing indirect searches to locate the platform. A similar proportion of trials were indirect searches over days 2-8 of training. Mice increasingly displayed directed searches, focal searches and direct paths such that, by the end of training, search was spatially specific on over 80% of trials. There were no major sex differences in strategy. The usefulness of strategy analyses (at least with default settings) for long probe trials is limited since spatially-specific strategies rely on IPE, which rapidly increases with trial duration. Additionally, animals will change strategies as they learn that the escape platform is not available in the expected location. Indeed, when the probe trial analysis was restricted to the first 10s, mice displayed focal and directed search strategies, indicating perseveration at the former platform location. When the analysis was conducted on longer segments, chaining was common, indicating that mice adopted a procedural strategy of searching in similar regions throughout the pool. Finally, when examining the entire probe trial, scanning and random searches dominated, indicating that mice eventually abandoned strategies that were no longer successful. During reversal, spatial specificity was initially very poor; mice primarily scanned, indicating preserved knowledge of the procedural requirements but no knowledge of the platform location. By the end of day 2 mice displayed levels of spatially-specific search strategies that were comparable to those at the end of the acquisition phase. Using the “add goal” feature, we also analyzed reversal strategies with respect to the original goal location Fig. 4b). This revealed a number of direct paths to the goal on the first day that quickly dissipated with additional trials as mice learning the new platform location.

**Figure 4:**
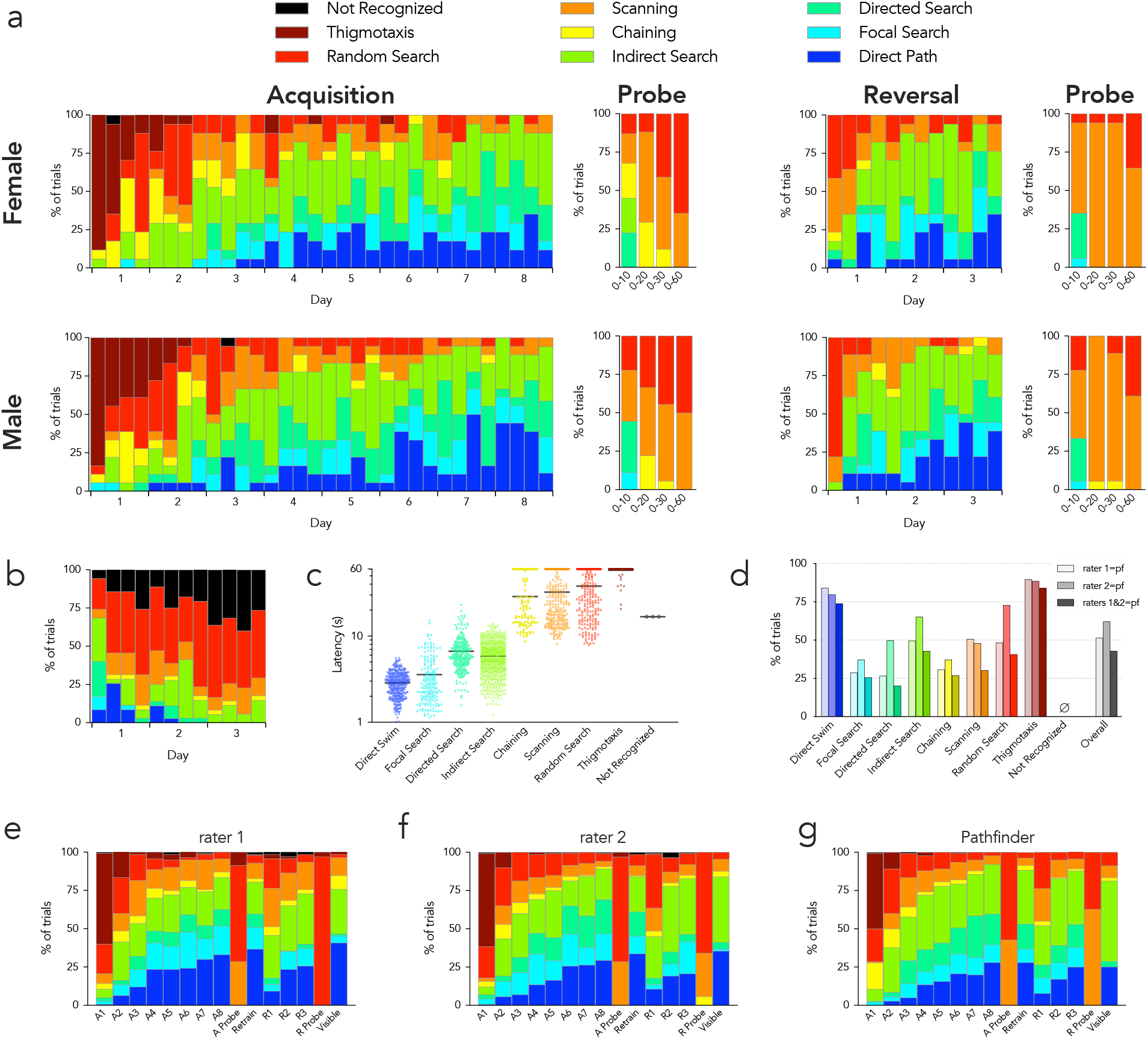
Pathfinder search strategy categorization of water maze performance. a) Search strategies for male and female mice. Each set of stacked bars indicates strategies used for the 4 acquisition and reversal trials for each day. Probe strategies are shown for the entire trial (0-60s) and for the first 10, 20 and 30s. b) Reversal strategies relative to the original platform location (indirect search excluded from analyses, since short swims that bypass the old location but quickly go to the new location become incorrectly classified as indirect searches with current settings). c) Escape latencies for all 1888 trials varied by strategy. Symbols indicate individual trials, bars indicate means (Kruskal Wallis test, P<0.0001; Dunn’s tests: direct path vs all others except focal search, P<0.0001; focal search vs all others except direct path, P<0.0001; directed search vs all others except indirect search, P<0.0001; indirect search vs all others except directed search, P<0.0001; chaining vs all others except scanning and random, P<0.01; scanning vs all others except chaining and thigmotaxis, P<0.01; random vs all others except chaining, P<0.05; thigmotaxis vs all others except random search, P<0.05). d) Manual vs automatic categorization. For each strategy assigned by Pathfinder, the proportion that received the same classification (manually) by 2 raters is shown. “Raters 1+2” indicates the percentage of Pathfinder classified-trials that also received the same classification by both raters. e) Search strategy classification by rater 1 for each day of testing. f) Search strategy classification by rater 2 for each day of testing. g) Search strategy classification by Pathfinder for each day of testing.

To investigate possible relationships between strategy and conventional measures of water maze performance, we examined escape latencies for each strategy type, over all trials (1888 trials from all 15 days of testing; Fig. 4c). Direct swim trials had the lowest latencies (2.9s on average) and was followed by the other spatially-specific strategies (focal search, 3.6s; directed search, 6.7s; indirect search, 5.8s). Non-specific strategies that avoided the pool wall were all significantly worse than the spatially specific strategies (chaining, 29s; scanning, 32s; random, 38s), and thigmotaxic trials were significantly worse than all other trial types (58s).

To determine how Pathfinder compared to subjective assessment of strategy, we compared Pathfinder categorizarion to manual scores generated by 2 independent raters (all trials). Rater 1 had experience in mouse behavior testing, but only brief training on water maze strategy classification. Rater 2 developed Pathfinder and had extensive experience with strategy classification. Figure 4d shows the proportion of Pathfinder-categorized trials that received the same strategy classification via the manual raters. The greatest correspondence between automatic and manual categorization was seen for direct swims and thigmotaxis (~80% for both). Automatic-manual consistency was much lower for the other strategies, ranged from 25-75% and differed for the 2 raters. Overall consistency between the 2 manual raters was 65%. These data highlight the difficulty of intuitively differentiating complex search paths. Interestingly, when we averaged strategy analyses over all 15 days of testing, automatic and manual categorization resulted in similar patterns (Fig. 4e-g). Thus, manual scoring is unreliable at the level of an individual trial, and human error can be masked when data are averaged.

To provide an intuitive visual inspection of search performance, we used Pathfinder to generate heatmaps of spatial occupancy at stages of testing that differed in spatial search patterns (Fig. 5). Averaged over all trials and across sexes, mice swam in close proximity to the pool wall on day 1 of initial acquisition. By days 3 and 8 search was increasingly focussed near the goal. Spatial preference was clearest on the probe trial, since these trials provided a longer temporal window to accumulate spatial occupancy samples. Day 1 of reversal testing resembled the probe trial, since mice spent the majority of time in the former platform location. By day 3, and on the probe trial, their spatial preference had shifted to the new, correct location. One set of heatmaps are presented using Pathfinder’s auto scale feature, which maximizes the color range within a trial and can be useful for visualizing within-trial details since it avoids saturation. However, by differentially scaling, it can also obscure or inflate differences across trials. We therefore include a second set of heatmaps that are all scaled equivalently.

**Figure 5:**
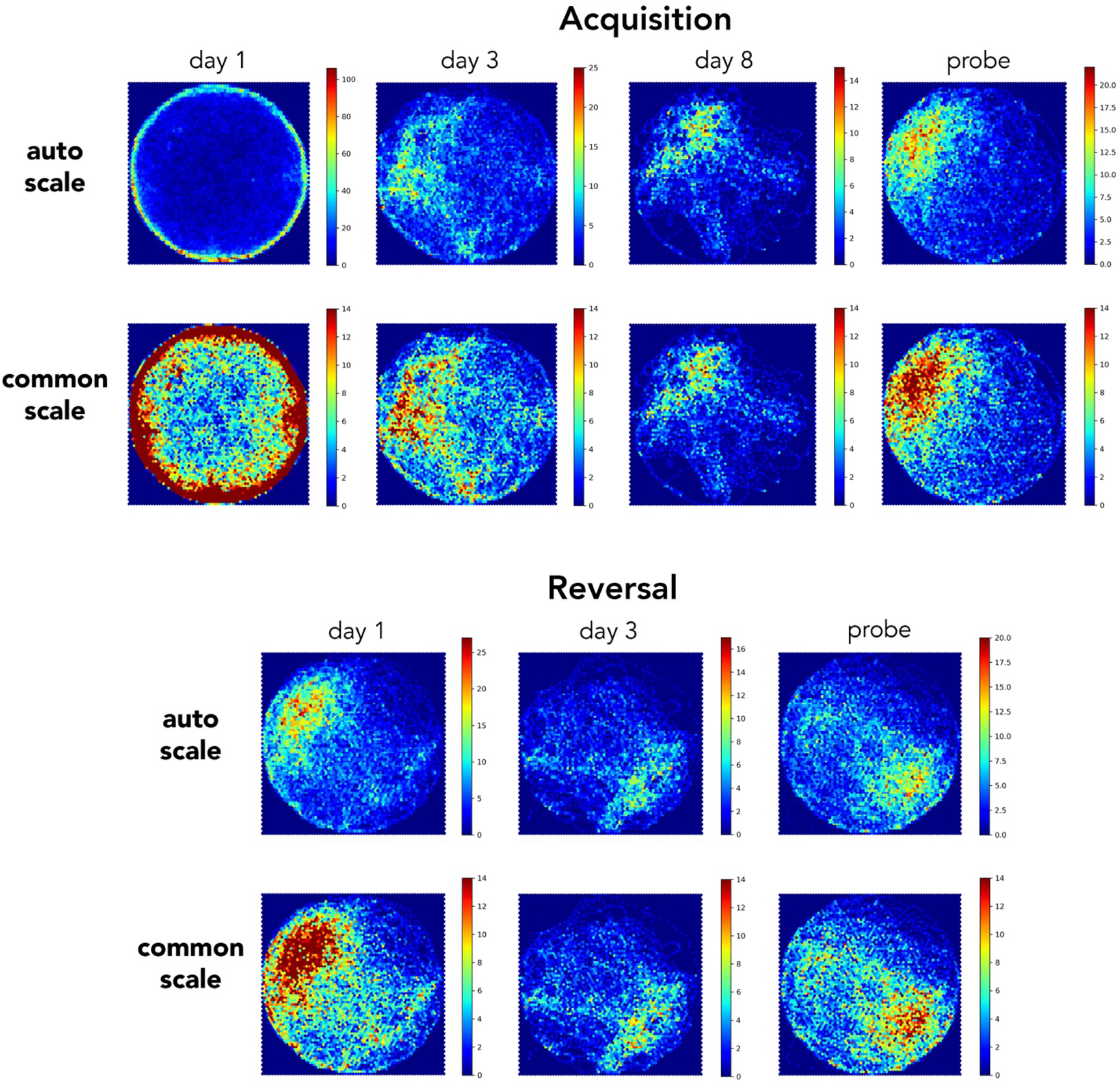
Heatmap visualization of spatial occupancy. Examples of heatmaps for various testing days (all trials from both sexes combined). Top rows: heatmaps were automatically scaled by Pathfinder, to occupy the full color spectrum and facilitate visualization of spatial occupancy within a given day. Bottom rows: heatmaps were set to a common scale, to facilitate comparison across days. Scale indicates number of samples within a spatial bin.

## DISCUSSION

Here we describe Pathfinder, an easy-to-use software package for analyzing patterns of spatial navigation. Pathfinder performs automatic classification of multiple search strategies that have been previously described in the rodent water maze, but it can also be used for analyzing navigational behavior in dry mazes, virtual mazes or any other environment where xy coordinates are provided. Currently, Pathfinder accepts inputs from 3 commonly-used, commercially-available tracking programs (Ethovision, Anymaze, Watermaze) and also the freely-available tracking software, ezTrack^26^. It requires no programming knowledge, but is open source and can be expanded by developers in the future. Using a mouse water maze dataset, we validated Pathfinder’s performance and found that mice progressed through a series of search strategies that had increasing levels of spatial search specificity, consistent with earlier reports^20,21 16–18,25,31^. Mice initially displayed thigmotaxic, random and chaining search strategies as they learned the procedural components of the task. Pathfinder effectively demonstrated that mice transitioned to spatially-specific, presumably hippocampal-dependent, strategies during the later stages of training. Pathfinder also revealed the reverse transition from spatial to procedural to random strategies in the probe trial. By analyzing reversal performance with respect to multiple goal locations, Pathfinder showed that mice redirect their spatial search from the previously-reinforced platform location to the new location. Mice displayed a variety of search strategies on any given day, even after escape latency performance had plateaued. Since manual classification based on static images of swim paths was slow and inconsistent, Pathfinder may therefore be a useful tool for objectively characterizing swim strategies in the rodent water maze and 2D spatial navigation in other behavioral paradigms.

The water maze was initially described nearly 40 years ago and quickly became popular due to the ease of training, strong motivation for escape, and consistent reliance on hippocampal function^3,32^. Escape latency and path length were quickly adopted as the primary measures of learning and, due to their simplicity and sufficiency for many experimental situations, they remain the most commonly-used metrics. However, they cannot always differentiate between behaviors that vary in the degree of spatial bias. For example, animals that employ a chaining strategy search nonspecifically but in some cases can reach the platform as fast as animals that perform a directed spatial search (Fig. 4c). Latency and path length are also less capable of detecting age-related impairments in spatial learning, prompting development of measures of proximity to the goal location, which has proven to be highly sensitive to group differences in both training and probe trial performance^9,27,33^. Our IPE proximity measure is similar to previous proximity measures with the exception that the cumulative ideal path distance is subtracted from the cumulative actual path distance to generate a path error measure. Finally, another recent metric that has been reported to be even more sensitive to group differences in spatial probe trial performance is entropy^28^. Entropy is originally a measure of thermodynamic disorder in a system but, when applied to the distribution of sampled sites in a maze, can also be used to measure the transition from high to low disorder in navigation, as animals focus their search on the precise goal location. By default, Pathfinder applies equal weighting to the path and goal components of the entropy measure and, compared to standard metrics, entropy was slightly better at detecting performance changes during the later stages of water maze acquisition.

Despite their convenience, even the most precise individual measures cannot distinguish between multiple possible strategies that an animal might employ to reach a goal location. Thus, strategy analyses may be valuable for identifying the role that different circuits play in guiding behavior. Consistent with the lower spatial resolution of ventral hippocampal place cells^34^, strategy analyses have found that the ventral hippocampus is particularly important for developing coarse, non-specific search patterns in the water maze and that increasingly spatially localized search depends on sequential recruitment of intermediate and then dorsal hippocampus^16^. Adult-born neurons are believed to promote memory precision and, indeed, blocking neurogenesis greatly reduced the adoption of spatially-specific search strategies^20^. Strategy analyses in animals have revealed spatial precision-related deficits in models of aging^25^, stroke^35^, traumatic brain injury^22,24^, autism^23^ and Alzheimer’s pathology^21,22^. With the advent of virtual reality, it has also become possible to test whether rodent water maze findings generalize to humans^15^. Indeed, hippocampal damage and CA1-specific lesions impair human water maze performance according to standard measures such as latency to reach the platform^36,37^. Human water maze experiments have also revealed superior spatial memory and greater spatial strategy use in younger individuals, and in males compared to females^14^. Here, we did not observe any sex differences, possibly because these mice had been previously tested in a Barnes maze^29^, which may have normalized stress reactivity and resulted in equal performance in males and females^38–40^.

Given the apparent utility of strategy classification, the question arises as to why it has not been used more extensively. One likely explanation is that it is not a standard feature of commercially-available software packages, therefore requiring time and programming experience to execute. Groups that have performed strategy analyses have developed their own software, using either a predefined parameter-based approach, like ours, or machine learning algorithms that classify based on user input^8,11,12,15–20,25,41^. Since most previous approaches have not been developed into freely-available software packages, Pathfinder may enable more widespread adoption of strategy analyses. Moreover, in conjunction with freely-available tracking programs such as ezTrack (which is already supported) or others^41^, users should be able to easily perform advanced navigation analyses at little cost.

It is worth noting that, with respect to water maze analyses, some behaviors (e.g. chaining and thigmotaxis) have been relatively well-described. In contrast, differences between spatially-specific search patterns (direct swim, directed search, focal search, indirect search) may be intuitive and quantifiable but the extent to which they are meaningful and result from distinct neural processes is less clear. Certainly, the fact that search strategies can now be easily quantified opens the door to future studies of the biology of complex navigation strategies. However, to some extent, strategy definitions are arbitrary and it is therefore incumbent upon the user to determine which behaviors are relevant for their experimental paradigm.

### Future developments and additional uses

One area where Pathfinder could be useful is for assessing spatial bias and choice behavior when there are multiple goal locations. Indeed, the water maze has been effectively used to study visuospatial goal discrimination^42,43^ and cue vs place-related choice behavior^44,45^. We have recently used Pathfinder to show that neurogenesis promotes spatial platform preference in a spatial alternation water maze, which was detected by a greater number of direct swims when the platform was in the rat’s preferred location than when it was in the non-preferred location^46^. Neurogenesis-deficient rats often vacillated between the two platform locations, similar to vicarious trial and error behavior that has been described at choice points in dry mazes^47^. Future software developments could possibly incorporate these types of movements between competing goal options to detect indecisiveness as animals refine goal-directed navigation behavior. Swim speeds are also currently not factored into Pathfinder’s classification scheme, but could provide useful additional information for strategies that incorporate goal expectancy^48,49^ or a transition between place- and cue-directed navigation^5^. In the water maze, multiple platform locations (> 2) are typically only used in matching-to-place variants where it is expected that subjects quickly forget previous goal locations^30^. However, there is evidence that search patterns can reflect memory for many individual goal locations as well as the overall distribution of goals, at recent and remote post-training intervals, respectively^6^. Since Pathfinder can analyze navigation with respect to an unlimited number of goal locations, it may be useful for future investigations of how multiple spatial goals interact to guide search.

Spatial navigation and exploration have been studied in many paradigms and so it is worth reiterating that Pathfinder could be applied to study navigation by any species, in any open 2D environment, and not just the water maze. For example, it could be used to measure the spatial precision of homing behavior^50,51^, spatial preferences of mammals or invertebrates in novel environments^52–54^, or navigation with respect to other environmental features that are known to drive firing of select populations of neurons, such as local and distal cues^55^, objects^56,57^ and environmental borders^58^. An array of virtual environments also opens the door to similar analyses of spatial navigation in humans^36,37,59,60^. Finally, eye tracking data, as humans and nonhuman primates explore 2D scenes, provides a measure of navigation that is analogous to rodent spatial exploration^61^. Indeed, hippocampal-damaged subjects display disorganized, inefficient search in a scene exploration task and are impaired according to several water maze-inspired metrics such as cumulative search error and heading angle error^62^. As a user-friendly application that can be further developed to accommodate differences between these various paradigms, Pathfinder may be a useful tool for characterizing complex spatial behavior and bridging findings across humans and animal models.

## ACKNOWLEDGEMENTS

The authors thank Sabri Snyder for contributing the name, Pathfinder, and Kurt Stover for contributing initial work towards automatic strategy classification. This work was funded by the Canadian Institutes of Health Research (JSS), the Natural Sciences and Engineering Research Council (JSS, REB), and the Michael Smith Foundation for Health Research (JSS, TPO).

## DATA AVAILABILITY

The datasets analysed in the current study are available from the corresponding author on reasonable request. The software, Pathfinder, is freely available at github.com/MatthewBCooke/Pathfinder.

